# Interaction of diabetes and smoking on stroke: A population-based cross-sectional survey in China

**DOI:** 10.1101/277293

**Authors:** Heqing Lou, Zongmei Dong, Pan Zhang, Xiaoping Shao, Ting Li, Chunyan Zhao, Xunbao Zhang, Peian Lou

## Abstract

**Objectives:** Diabetes and smoking are known independent risk factors for stroke; however, their interaction concerning stroke is less clear. We aimed to explore such interaction and its influence on stroke in Chinese adults.

**Design:** Cross-sectional study.

**Setting:** Community-based investigation in Xuzhou, China.

**Participants:** A total of 39,887 Chinese adults who fulfilled the inclusion criteria were included.

**Methods:** Participants were selected using a multi-stage stratified cluster method, and completed self-reported questionnaires on stroke and smoking. Type 2 diabetes mellitus (DM2) was assessed by fasting blood glucose or use of antidiabetic medication. Interaction, relative excess risk owing to interaction (RERI), attributable proportion (AP), and synergy index (S) were evaluated using a logistic regression model.

**Results:** After adjustment for age, sex, marital status, educational level, occupation, physical activity, body mass index, hypertension, family history of stroke, alcohol use, and blood lipids, the relationships between DM2 and stroke, and between smoking and stroke, were still significant: odds ratios were 2.75 (95% confidence interval [CI]: 2.03–3.73) and 1.70 (95% CI: 1.38–2.10), respectively. In subjects with DM2 who smoked, the RERI, AP, and S values (and 95% CIs) were 1.80 (1.24–3.83), 0.52 (0.37–0.73), and 1.50 (1.18–1.84), respectively.

**Conclusions:** The results suggest there are additive interactions between DM2 and smoking and that these affect stroke in Chinese adults.

**Article Summary: Strengths and limitations of this study:**

- The strengths of this study were that a large sample population was randomly selected from the general population of Xuzhou and many confounding risk factors were adjusted for.
- Owing to the cross-sectional design, we could not determine a causal combined relationship among diabetes, smoking and stroke.
- We were not able to control for some important and well-known risk factors of diabetes, such as heart rate and cardiovascular causes.
- We did not measure fresh fruit consumption, which is causally related to stroke.

## INTRODUCTION

Stroke is an ongoing global health problem. In 2016, there were 5.53 million from stroke worldwide. Stroke was also the second most common cause of premature mortality and secondary disability.^1^ The 2015 Report on Chinese Stroke Prevention indicates stroke as the leading cause of death in China, with incidence increasing by 8.7% per annum.^2^ Stroke rates in China are higher than those in Western and other Asian countries.^3,4^ Moreover, the number of stroke patients in China is likely to increase because of lifestyles, demographic changes, and inadequate control of major risk factors for stroke.^2^ It is therefore important to identify and prevent these risk factors.

The major risk factors for stroke are hypertension, diabetes mellitus, hypercholesterolemia, smoking, physical inactivity, obesity, and a family history of stroke.^5-8^ Although these play a role in the development of stroke, its formation is not entirely caused by a single risk factor. The more risk factors a person has, the greater the likelihood of incurring a stroke.^7^ In fact, >90% of the global burden of stroke in 2013 was attributable to the combined effect of all modifiable risk factors.^8^

People with comorbid diabetes and smoking may represent a subgroup with high risk of developing stroke; however, few studies have examined the interaction of diabetes and smoking with regard to stroke. The primary aim of this cross-sectional study was to examine the interaction of type 2 diabetes mellitus (DM2) and smoking on stroke in Chinese adults. A secondary aim was to evaluate the associations between DM2 and stroke, and between smoking and stroke.

## MATERIALS AND METHODS

### Study design and recruitment criteria

This population-based, cross-sectional survey was conducted in Xuzhou City, Jiangsu Province, China, from February to June 2013. The sample was selected with two-stage probability proportional to size, from all 11 regions in Xuzhou. In the first stage, five subdistricts/townships in urban/rural areas were selected in accordance with the population of each subdistrict/township from each region, with probability proportional to size sampling. In the second stage, five communities/villages were selected in accordance with the population of each community/village from each subdistrict/township with probability proportional to size sampling. In the final stage, one person ≥18 years old and who had lived in his or her current residence for ≥5 years was selected from each household through use of a Kish selection table. Those who met either or both of the following criteria were excluded: (1) previous diagnosis of neuropathy, psychosis, or unclear speech or (2) member of the floating population or temporary residents. A total of ≥13,500 people were selected, assuming an estimation incidence of stroke of 2.0%,^9^ with 90% power, α=0.05, and allowing for a dropout rate of 15%. Trained interviewers interviewed participants face-to-face on the day of the participants’ regular medical appointments. All participants underwent 12-h overnight fasting and blood sampling to test basic fasting plasma glucose. After blood sampling, each received a health examination and completed a structured questionnaire inquiring on demographic information, medical history, medication history, and smoking, alcohol consumption, and exercise habits.

### Key measurements

Stroke was assessed using subjects’ self-reported responses, and defined as an acute disturbance of focal areas in the brain lasting for ≥24 h and thought to be caused by intracranial hemorrhage or ischemia.^10^ We examined the medical records of participants reporting a diagnosis of stroke to check that participants satisfied this definition. The diagnosis was also confirmed through computed tomography and magnetic resonance imaging scans. Detailed clinical information about stroke was based on the International Classification of Disease, 10th Revision, codes I60–I64. DM2 was defined as fasting blood glucose ≥7.0 mmol/L, any use of antidiabetic medication, or self-reported history of DM2.^11^ Hypertension was defined as systolic blood pressure ≥140 mmHg or diastolic blood pressure ≥90 mmHg, any use of antihypertensive medication, or self-reported history of hypertension.^12^

### Covariates

Age, sex, current employment status, marital status, level of education, cigarette smoking, alcohol consumption, physical activity, and family history of diseases, including DM2, hypertension, and stroke, were assessed using a standardized questionnaire. Employment status was categorized as manual, non-manual, unemployed, or retired. Education was categorized as below high school, high school, or above high school. Lifestyle variables included cigarette smoking, alcohol consumption, and physical activity level. Cigarette smoking was defined as having smoked at least 100 cigarettes in one’s lifetime. Information was obtained on the amount and type of alcohol consumed during the previous year, and alcohol drinking was defined as consumption of ≥30 g per week for ≥1 year. Regular leisure-time physical activity was defined as participation in moderate or vigorous activity for ≤30 min per day, ≥3 days a week. Each participant’s height (to the nearest 0.1 cm) and weight (to the nearest 0.1 kg) in light indoor clothing were measured. Body mass index (BMI; in kg/m^2^) was calculated; categorized as underweight (<18.5 kg/m^2^), normal weight (18.5-24.0 kg/m^2^), or overweight/obese (>24.0 kg/m^2^).^13^Dyslipidemia was defined as use of any lipid-lowering medication or self-reported history of the condition.

### Statistical analysis

Participants were divided into four groups in accordance with their smoking status and DM2: non-smokers with no DM2, smokers with no DM2, non-smokers with DM2, and smokers with DM2. Statistical analysis was performed using IBM SPSS for Windows, Version 16.0 (SPSS Inc., Chicago, IL, USA). The general characteristics of continuous variables were compared across the four subgroups using analysis of variance. The categorical variables were expressed as a percentage and the groups were compared using a chi-squared test. Logistic regression analysis was performed to estimate the probability of having a stroke and 95% confidence interval (CI) for each risk factor category stratified by DM2 and smoking, adjusting for age, sex, occupation, education, marital status, BMI, physical activity, drinking status, hypertension status, and family history of disease, including DM2, hypertension, and stroke.

Biological interactions should be based on an additive scale rather than a multiplication scale.^14,15^ We therefore used three measures to estimate biological interactions between DM2 and smoking: relative excess risk owing to interaction (RERI), attributable proportion (AP) owing to interaction, and synergy index (S). RERI is the excess risk attributed to interaction relative to the risk without exposure to diabetes and smoking. AP refers to the attributable proportion of disease caused by interaction in subjects with exposure to both variables. S is the excess risk from exposure to both variables when there is a biological interaction relative to the risk from exposure to both variables without interaction. In the absence of additive interactions, RERI and AP equal 0.^14,16^ In the current study, RERI >0, AP >0, and S >0 indicated statistical significance, set at p <0.05 (two-tailed).

### Ethics approval and consent to participate

The study protocol was approved by Xuzhou Center for Disease Control and Prevention. The procedures followed were in accordance with the standards of the ethics committee of the Xuzhou Center for Disease Control and Prevention and with the Declaration of Helsinki (1975, revised 2000). Written informed consent was obtained from all participants.

## RESULTS

### General characteristics of participants

Of the 41,658 individuals initially sampled, 1253 did not respond to the smoking question or complete the blood glucose tests, and 518 did not meet our study criteria. A final total of 39,887 adults (19,178 men and 20,709 women) with complete data were included in the analysis (response rate: 95.7%). Table 1 shows the characteristics of the study population. The proportion of participants with DM2 was 7.00% (2783/39,887), of stroke subjects with DM2 was 16.67%, and of non-stroke subjects with DM2 was only 6.75%; there was a statistically significant difference between the two groups (χ^2^ = 135.92, p <0.001). The proportion of smokers was 20.26% (8082/39,887), stroke participants who smoked was 32.24%, and non-stroke participants who smoked was 19.98%; there was a statistically significant difference in smoking between stroke and non-stroke patients (χ^2^ = 83.49, p <0.001).

**Table 1.**
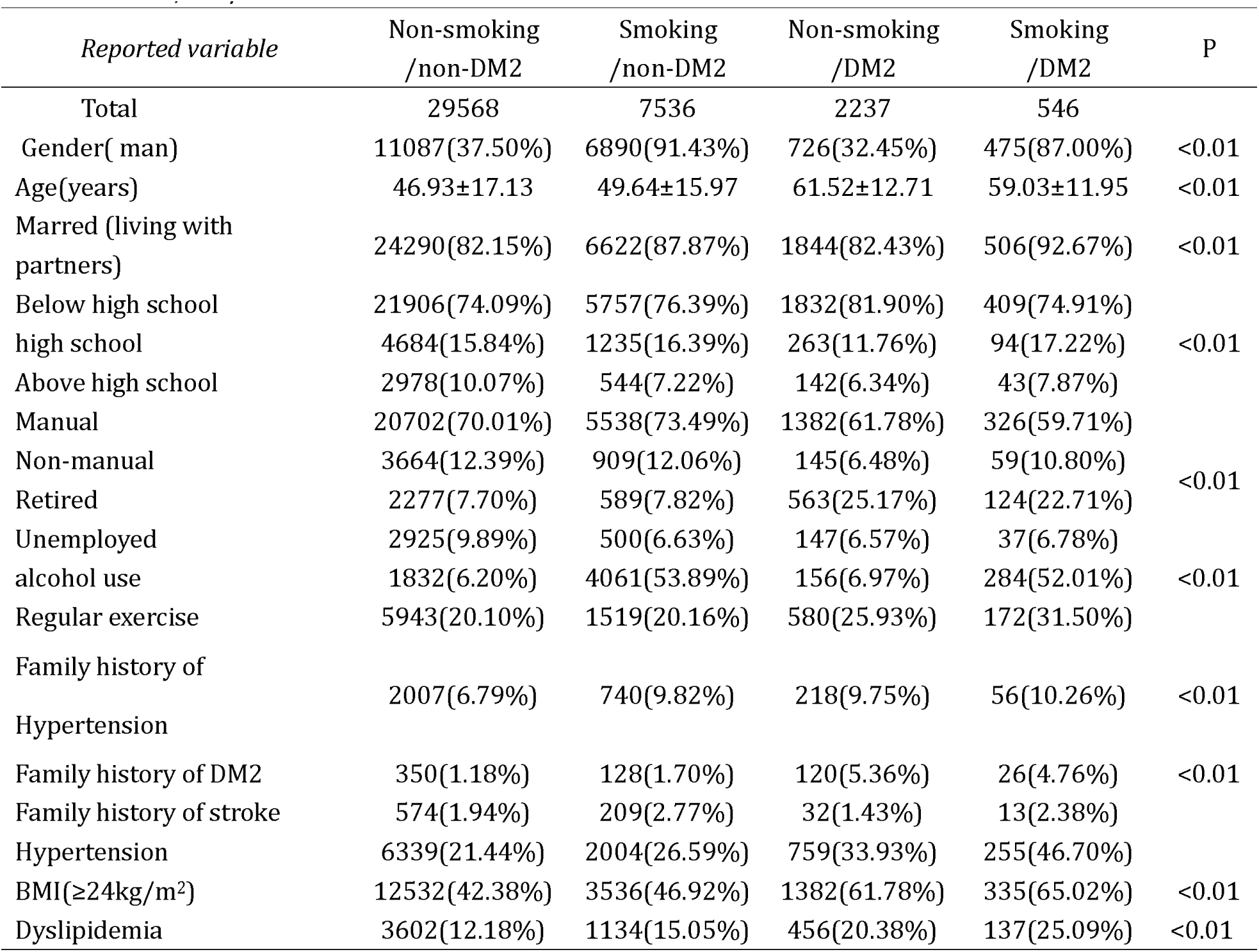
General characteristics of diabetes and smoking in the study population (n = 39,887)

### Association of diabetes and smoking with stroke

The 5.50% incidence of stroke in subjects with DM2 exceeded the 2.06% incidence in those with no DM2 (χ^2^ = 139.11, p <0.001). Individuals who smoked had a higher stroke incidence compared with non-smokers (3.36% vs. 1.96%; χ^2^ = 83.49, p <0.001; see Table 2). Subjects with DM2 had a significantly increased risk of stroke compared with those with no DM2 (OR: 2.65, 95% CI: 1.70–4.41, p < 0.001), after adjusting for confounders (see Table 3). The risks of ischemic and hemorrhagic stroke were increased by DM2, after adjusting for confounders; the ORs were 2.71 (95% CI: 1.72–4.49) and 1.82 (95% CI: 1.34–3.35), respectively. Smokers had a significantly increased risk of stroke compared with non-smokers (OR: 1.83, 95% CI: 1.59–2.14, p <0.001), after adjusting for confounders (see Table 3). The risks of ischemic and hemorrhagic stroke were increased by smoking, after adjusting for confounders; the ORs were 1.32 (95% CI: 1.12–2.53) and 1.95 (95% CI: 1.40–3.41), respectively.

**Table 2.** Associations between smoking, diabetes, and stroke

| Variables |  | Stroke | Non-stroke | Unadjusted OR(95%CI) | Adjusted OR (95%CI) | P |
| --- | --- | --- | --- | --- | --- | --- |
| Smoking | No | 622(67.76%) | 31183(80.02%) | 1.91 | 1.83 | <0.01 |
|  | Yes | 296(32.24%) | 7786(19.98%) | (1.63-2.31) | ( 1.59-2.14 ) |  |
| DM2 | No | 765(83.33%) | 36339(93.25%) | 2.76 | 2.65 | <0.01 |
|  | Yes | 153(16.67%) | 2630(6.75%) | (1.77-4.68) | ( 1.70-4.41 ) |  |

Adjusted for age, sex, education, marriage status, employment, family history of diabetes, family history of hypertension, family history of stroke, alcohol consumption, physical activity, hypertension, diabetes or smoking, body mass index, and lipids.

**Table 3.**
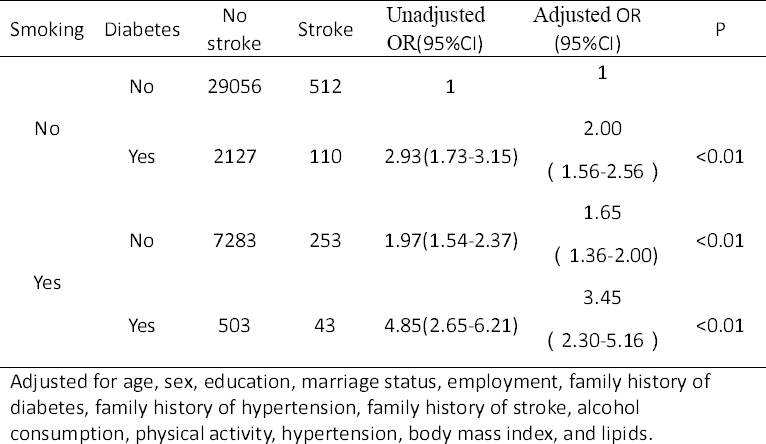
Odds ratios (ORs) for the association between smoking and stroke by diabetes among participants

### Interaction between diabetes and smoking in relation to stroke

Individuals who only had DM2 or only smoked had a significantly increased risk of stroke compared with those who did not have DM2 and did not smoke (OR: 2.00, 95% CI: 1.56-2.56; OR: 1.65, 95% CI: 1.36-2.00; respectively; all p <0.001), after adjusting for confounders. Table 3 shows the results from the multiple logistic regression models. The incidence of stroke was greatest in those who had DM2 and smoked (OR: 3.45, 95% CI: 2.30-5.16, p <0.001), after adjusting for confounders.

### Sensitivity analysis

There was a strong additive interaction between DM2 and smoking (RERI: 1.80, 95% CI: 1.24–2.14; AP: 0.52, 95% CI: 0.37–0.73; and S: 1.50, 95% CI: 1.18–1.84, respectively); 52% of stroke occurrence was attributed to the interaction between DM2 and smoking (Table 4).

**Table 4.** Measures for estimating biological interaction between smoking and diabetes for prevalence of stroke in participants

| Measures of biological interaction | Estimate (95% CI) |
| --- | --- |
| RERI | 1.80(1.24 - 3.84) |
| AP | 0.52(0.37 - 0.73) |
| S | 1.50(1.18 - 1.84) |

Reference group is no smoking with non-diabetes.
Adjusted for age, sex, education, marriage status, employment, family history of diabetes, family history of hypertension, family history of stroke, alcohol consumption, physical activity, hypertension, body mass index, and lipids.

## DISCUSSION

There were two main findings in this study. First, there was an interaction of DM2 and smoking with relation to the incidence of stroke. Second, DM2 and smoking were found to be significantly associated with increased risk of stroke in Chinese adults, independent of potential confounders such as age, sex, occupation, education, marital status, BMI, physical activity, drinking status, hypertension status, and family history of diseases including diabetes, hypertension, or stroke.

The proportion of participants with DM2 in this study was 19.67%, which is lower than that reported in the China National Stroke Registry Study (27.0%),^17^ lower than that reported in a comparison of risk factors for ischemic stroke in Chinese versus Caucasian individuals (25.0%),^18^ and higher than that reported in a study of the Chinese National Health Insurance program (3.8%)^19^ and a review of stroke in China (4%–15%).^20^ The present results are also inconsistent with figures for diabetes in stroke patients reported for other countries.^21,22^ This discrepancy may be due to differences in study design, sampling size, region, diabetes diagnosis, and the clinical features of stroke, but could also be a result of ethnic group differences, as many studies have shown that stroke incidence rates differ among ethnic groups.^23^

Numerous epidemiologic studies, including cross-sectional studies and prospective cohort studies, have demonstrated associations between diabetes and stroke.^22,24-27^ A population-based study of 173,979 discharged patients admitted with hemorrhagic stroke was conducted in Spain from 2003 to 2012. The authors of that study reported a positive association between diabetes and stroke (incidence rate ratio: 1.38, 95% CI: 1.35–1.40 for men; incidence rate ratio: 1.31, 95% CI: 1.29–1.34 for women).^22^ Folsom et al. reported that the relative risk (RR) of ischemic stroke was 2.22 (95% CI: 1.5–3.2) for individuals with diabetes after adjustment for other stroke risk factors in a 6–8-year follow-up.^24^ Iso et al.^25^ reported that the association between non-embolic ischemic stroke and diabetes was particularly strong among non-hypertensive subjects with higher subscapular skinfold thickness values; the multivariate RR was 4.9 (95% CI: 2.5–9.5) in a 17-year prospective cohort study of 10,582 Japanese individuals (4287 men and 6295 women) aged 40–69 years. A systematic review and meta-analysis of 64 cohort studies with 775,385 individuals showed that diabetes is consistently associated with increased risk of stroke; the pooled maximum-adjusted RR of stroke associated with diabetes was 2.28 (95% CI: 1.93–2.69) for women and 1.83 (95% CI: 1.60–2.08) for men.^26^ Liao et al. also reported that diabetic patients have an increased risk of stroke (adjusted hazard ratio: 1.75, 95% CI: 1.64–1.86) compared with those without diabetes; associations between diabetes and stroke risk were significant for both sexes and all age groups.^27^ The present findings also demonstrated an association between diabetes and stroke.

Robson et al.^28^ confirmed that poor blood sugar control increases the risk of stroke. One prospective cohort study of 467,508 men and women aged 30–79 years with no history of diabetes or stroke showed that each 1-mmol/L (18-mg/dL) higher than usual increase of random plasma glucose level above 5.9 mmol/L (106 mg/dL) was associated with ischemic stroke (adjusted hazard ratio: 1.08, 95% CI: 1.07–1.09).^29^ Moreover, a study of 4669 patients who had incurred a minor stroke revealed during a 3-month follow-up that stroke patients with diabetes experienced stroke recurrence and disability.^30^ Therefore, effective glycemic control not only reduces the incidence of stroke but also can reduce stroke recurrence and associated disability.

A higher proportion of stroke than non-stroke patients tend to be smokers. Indeed, Wang et al. reported that 48% of stroke patients smoked.^31^ Tsai et al.^18^ reported a figure of 38% and our results showed 32.24%. Although the proportions differ, they are all quite high. The discrepancy may reflect a bias in the reporting of smoking status among study participants.

Many studies have shown that smoking is a strong risk factor for development of stroke.^32-34^ The British Regional Heart Study, which included 7735 men aged 40–59 years, showed that after full adjustment for other risk factors, current smokers had a nearly fourfold RR of stroke compared with never-smokers (RR: 3.7, 95% CI: 2.0–6.9). Ex-cigarette smokers showed lower risk than current smokers, but showed excess risk compared with never-smokers (RR: 1.7, 95% CI: 0.9–3.3, p = 0.11).^33^ During a mean follow-up of 8.5 years, the risk for all forms of stroke significantly increased (RR: 0.9–3.3, 95% CI: 1.4–24) and increased for ischemic strokes (RR: 4.8, 95% CI: 1.2–20) among cigarette-smoking women with a cigarette-smoking spouse compared with those with a non-smoking spouse and after adjusting for other cardiovascular risk factors.^34^ However, a systematic review and meta-analysis of 81 cohorts in Asia, including 3,980,359 individuals and 42,401 strokes, showed smoking did not contribute to the risk of stroke.^35^ The proportion of Chinese stroke patients who smoke is higher than that of Caucasians.^18^ One systematic review and meta-analysis of 15 cohort studies and 178 case-control studies found smoking was an independent risk factor for stroke (pooled RRs: 1.27, 95% CI: 1.21–1.35) in a Chinese population.^7^ More frequent smokers are more likely to incur a stroke.^36,37^ A meta-analysis that included 16,886 men and 18,539 women without known diabetes revealed hemoglobin A1c was 0.10% (95% CI: 0.08–0.12, 1.1 mmol/mol [0.9–1.3]) higher in current smokers and 0.03% (95% CI: 0.01–0.05, 0.3 mmol/mol [0.1–0.5]) higher in ex-smokers than in never-smokers.^38^ Therefore, our findings support previous evidence smoking is associated with stroke in Chinese populations.^4^ However, the present study only wanted to observe the interaction of smoking and diabetes on stroke, the cigarette smokers were not categorized as current, former and never smokers. Therefore, when compared the association between smoking and stroke of our study with that of others should be carefully.

One study reported that in comparison with nonsmoking patients with no diabetes mellitus or hypertension, patients with those conditions and who smoked had a threefold increase in prevalence of peripheral vascular disease and a 3.5-fold increase in cerebrovascular disease^[39]^. Our results reinforce such findings.

The pathophysiological mechanisms of hyperglycemia induce oxidative stress; promote formation of advanced glycosylation end products;^40,41^ increase blood–brain barrier permeability and inflammatory responses;^42^ lead to accumulation of reactive oxygen species/reactive nitrogen, inflammation, and mitochondrial dysfunction;^43^ lead to cellular dysfunction; damage vascular tissue; inhibit endogenous vascular protective factors; alter vascular homeostasis;^44^ raise levels of reactive oxygen species and advanced glycation end products; decrease levels of mitochondrial superoxide dismutase;^41^ and correlate with endothelial cell dysfunction and nitric oxide production.^45^All these actions contribute to accelerating the atherosclerotic process. Therefore, diabetic patients are more prone to incurring stroke.

Cigarette smoking is associated with increased reactive oxygen species, oxidative stress, blood–brain barrier permeability, sympathetic activation and nitric oxide production, reduced cerebral blood flow and serum superoxide dismutase levels, attenuation of the vasodilation of cerebral arterioles, and induction of atherosclerosis and thrombosis.^46-49^ Moreover, cigarette smoke elevates serum levels of advanced glycation end products and reduces soluble receptors for those end products, resulting in development of atherosclerosis and related stroke.^50-52^ Smoking is therefore correlated with increased risk of stroke.

Collectively, diabetes and smoking induce oxidative stress and nitric oxide production; increase reactive oxygen species, blood–brain barrier permeability, and the level of advanced glycation end products; and reduce cerebral blood flow and serum superoxide dismutase. Therefore, diabetic patients who smoke also have a greater risk of stroke.

The strengths of the current study are that we used a community-based multistage sampling design, large sample size, and randomly selected participants. However, the study has several limitations. First, because of the cross-sectional design, we could not determine a causal relationship between DM2, smoking, and stroke. Second, we were unable to control for some important and well-known risk factors of stroke, such as heart rate^53^ and cardiac causes.^6^ Third, we did not measure fresh fruit consumption, ^54^which is causally negative related to stroke. Finally, patients self-reported their cigarette smoking status; therefore, the risk of misclassification and recall bias with the definition of smoking could not be avoid.

## Conclusion

The results of this cross-sectional study indicate diabetic patients who smoke are 3.5 times more likely to develop stroke than non-diabetics who do not smoke. Diabetes and smoking had a combined positive correlation with stroke. These results have important public health implications. Among Chinese adults, the current rate of smoking is as high as 28.3% ^55^ and DM2 prevalence is 11.6%.^56^ Therefore, it is important to implement stroke-prevention measures aimed at reducing smoking and improving glycemic control in diabetic patients in China.

## COMPETING INTERESTS

The authors declare that they have no competing interests.

## Acknowledgments

We thank all the participants involved in the survey. We also thank the Regional Centers for Disease Control and Prevention as well as clinics in Xuzhou for their collaboration.

## Funding

This research was funded by the Preventive Medicine research projects of Jiangsu Province Health Department in 2015 (Y2015010) and the Science and Technology projects of Xuzhou City in 2015 (KC15SM046). The researchers were independent from the funders. The study funders had no influence on the study design, data collection, analysis, interpretation of data, writing of the report, or decision to submit the article for publication.

## Duality of interest

The authors declare there is no duality of interest associated with this manuscript.

## Authors’ contributions

HL wrote/edited the manuscript and created tables. ZD, PZ, XS, TL, CZ, XZ, and PL contributed to the discussion and reviewed/edited the manuscript. XZ conceptualized the study. PL is the guarantor of this work and, as such, had full access to all data in the study and takes responsibility for the integrity of the data and accuracy of the data analysis. All authors read and approved the final manuscript.

## Availability of data and materials

All data relevant to the given manuscript have been stored in a separate file that can be made freely available to external investigators upon request.

